# An Ohio State Scenic River Shows Elevated Antibiotic Resistance Genes, Including *Acinetobacter* Tetracycline and Macrolide Resistance, Downstream of Wastewater Treatment Plant Effluent

**DOI:** 10.1101/2021.04.26.441562

**Authors:** April Murphy, Daniel Barich, Siobhan Fennessy, Joan L. Slonczewski

## Abstract

The entry of antibiotic resistance genes (ARGs) into aquatic systems has been documented for large municipal wastewater treatment plants, but there is less study of the impact of smaller plants that are situated on small rural rivers. We sampled water metagenomes for ARG and taxa composition from the Kokosing River, a small rural river in Knox County, Ohio, which has been designated an Ohio State Scenic River for retention of natural character. Samples were obtained 1.0 km upstream, 120 m downstream, and 6.4 km downstream from the effluent release of the Mount Vernon wastewater treatment plant (WWTP). ARGS were identified in metagenomes using ShortBRED markers from the CARD database screened against UniPROT. Through all seasons, the metagenome just downstream of the WWTP effluent showed a substantial elevation of at least 15 different ARGs, including 6 ARGs commonly associated with *Acinetobacter baumannii* such as *msrE, mphE* (macrolide resistance) and *tet*(39) (tetracycline resistance). The ARGs most prevalent near the effluent pipe persisted 6.4 km downriver. Using MetaPhlAn2 clade-specific marker genes, the taxa distribution near the effluent showed elevation of reads annotated as *Acinetobacter* species as well as gut-associated taxa, Bacteroides and Firmicutes. The ARG levels and taxa prevalence showed little dependence on seasonal chlorination of the effluent. Nitrogen and phosphorus were elevated near the effluent pipe but had no consistent correlation with ARG levels. We show that in a rural river microbiome, year-round wastewater effluent substantially elevates ARGs including those associated with multidrug-resistant *A. baumanii*.

**IMPORTANCE:** Antibiotic resistance is a growing problem worldwide, with frequent transmission between pathogens and environmental organisms. Rural rivers can support high levels of recreational use by people unaware of inputs from treated wastewater, while WWTPs can generate a small but significant portion of flow volume into a river surrounded by forest and agriculture. There is little information on the rural impacts of WWTP effluent on the delivery and transport of antibiotic resistance genes. In our study, the river water proximal to wastewater effluent shows evidence for the influx of multidrug-resistant *Acinetobacter baumanii*, an opportunistic pathogen of concern for hospitals but also widespread in natural environments. Our work highlights the importance of wastewater effluent in management of environmental antibiotic resistance, even in high quality, rural river systems.

## INTRODUCTION

Environmental sources of antibiotic resistance increasingly threaten global public health (1–3). Antibiotics from clinical use and livestock husbandry can promote the development of resistant bacteria, and they readily pollute urban and rural waterways (4–6). Even very low concentrations of antimicrobial drugs select for resistance (7). Antibiotic resistance genes (ARGs) that enter environmental microbial communities have the potential for transfer to pathogenic bacteria (8). Yet the public is rarely aware of the potential for exposure to ARG-carrying organisms in rural aquatic systems, particularly those designated for preservation by government agencies such as the Ohio Scenic Rivers Program (ohiodnr.gov).

A major source of ARGs and antibiotics in aquatic systems is the effluent of wastewater treatment plants (WWTPs) (9–11). Wastewater treatment may actually select for increased antibiotic resistance of potential pathogens such as *Acinetobacter* species (12, 13). It is important to understand the potential of WWTP to transfer ARGs as well as resistant microbes into rural streams, where they may disturb autochthonous microbial communities and spread drug resistance to human microbiomes. We investigated the impact of WWTP effluent on the taxa distribution and ARG counts in the Kokosing River, a rural river designated as a state “Scenic” River by the Ohio Department of Natural Resources (ODNR) as well as meeting the criteria for Exceptional Habitat by the Ohio Environmental Protection Agency (Ohio EPA) due to its high species diversity and high ecological condition (14).

The river microbiome may be affected by WWTP effluent in various ways: by elevation of phosphorus, nitrogen, and organic nutrients; by introduction of exogenous microbes and antibiotics; and by introduction of DNA including ARGs. The WWTP in our study chlorinates effluent only during the months of May through October, so we compared both conditions. While chlorination effectively decreases bacterial biomass by three log units (15, 16), it does not fully remove ARGs from effluent. Some studies show partial decrease of ARGs by chlorine (17, 18), whereas others show that chlorination may increase the effluent content of ARGs and promote their conjugative transfer (19, 20). Various stress conditions in the WWTP can co-select antibiotic resistances and virulence properties (21). In some cases the release of heavy metals, antibiotics, and other compounds into receiving rivers further propagates resistance by selecting for ARGs that encode multidrug efflux pumps (10, 22–24).

The establishment of antibiotic resistance in environmental microbial communities can be controlled when municipalities reduce antibiotic use (25). Therefore, understanding the impact of ARG pollutants on rural river resistomes is important for understanding the lasting potential of resistance in the environment. River resistomes offer the opportunity for surveillance of opportunistic pathogens that move between environment and human host, such as the ESKAPE pathogen *Acinetobacter baumanii* (26–28). The ESKAPE acronym comprises six leading hospital-acquired pathogens with multidrug resistance (29). While *A. baumanii* is known for hospital transmission, recent reports indicate community acquisition of strains that carry ARGs on plasmids (30, 31). In the Kokosing River we examined evidence for *Acinetobacter* ARGs such as *tet(39)* (32, 33) and *msrE, mphE* (34).

To understand how WWTP effluent with secondary treatment might alter rural river microbial communities, we sampled sites upstream, just downstream, and further downstream of the effluent release of the Mount Vernon WWTP on the Kokosing River. The Kokosing river in east-central Ohio, USA, flows 92 km into the Walhonding River, a part of the watershed of the Mississippi River (35). The Kokosing is included in Ohio’s Scenic Rivers Program; “scenic” designates “a waterway that retains much of its natural character for the majority of its length” (ODNR, ohiodnr.gov). The river is designated for: Exceptional Warmwater Habitat, Agricultural Water Supply, Industrial Water Supply, and Primary Contact Recreation (14). The river is used regularly for recreation by the local residents, including students from an undergraduate college (approximately 1800 students) situated at the downstream site reported by this study.

Nevertheless, the Ohio EPA recognizes some localized impairment of the Kokosing’s warmwater habitat and use for recreational activities (mywaterway.epa.gov/). Our study focused on a segment of the Kokosing in Knox County, proximal to the WWTP that serves the City of Mount Vernon (pop. 17,000). Mount Vernon includes surrounding suburban and rural homes as well as a 65-bed hospital. The WWTP system diagram is presented in Figure S1. The design flow is 5.0 MGD; actual discharge rates vary from 2.4-16.0 MGD (36). During our study dates the discharge accounted for 2-7% of the river’s daily flow rate (Table S1, Supporting Material). This fraction is small compared to the base flow contribution of municipal WWTP effluent to some rivers (37). Because it represents a small proportion of the river discharge, we asked whether the WWTP effluent would affect the microbiome of the system downriver of the plant. Small wastewater plants are situated approximately 25 km upstream (Village of Fredericktown, design flow 0.70 MGD) and 8 km downstream (Village of Gambier, 0.45 MGD). All of these plants disinfect their effluent by chlorination during six months of the year (May 1st through October 31st).

We focussed our study on the river water microbiomes upstream, midstream (proximal to effluent pipe) and downstream of the Mount Vernon WWTP. We examined how ARG numbers are associated with the WWTP; and how much ARG elevation may persist downstream of the effluent.

## MATERIALS AND METHODS

### Water sampling and metadata

All water samples were obtained from the Kokosing River, Knox County, Ohio. Water samples were obtained at three sites on the river to yield data on water quality upstream of the WWTP effluent, just downstream of the WWTP in the mixing zone where wastewater is mixed with river water (Midstream site), and further downstream where plant effluent has been fully diluted (**Fig. 1**). The Upstream site (coordinates 40.38368, −82.47042) lies approximately 1.0 km upstream of the Mount Vernon WWTP effluent discharge. WWTP discharge rates and river flow rates on the dates of sample collection are presented in Table S1. The Midstream site, nearest the WWTP (40.378007, −82.467822) is located approximately 120 m downstream of the effluent release pipe, within the mixing zone of the plant, where the effluent is initially mixed with river water. The Downstream site (40.376038, −82.40346) lies approximately 6.4 km downstream of the WWTP. The next nearest site where wastewater enters the Kokosing is the Fredericktown WWTP, a small plant (0.70 MGD) approximately 25 km upstream of the Mount Vernon WWTP. The three river sites were sampled using identical procedures at six dates throughout the year: October 27, November 3, and December 3, 2019; and April 13, May 28, and June 25, 2020. The WWTP effluent undergoes chlorination before discharge only from May 1^st^ through October 31^st^; thus, only the October, May and June samples occurred during the time that effluent was chlorinated.

**Figure 1.**
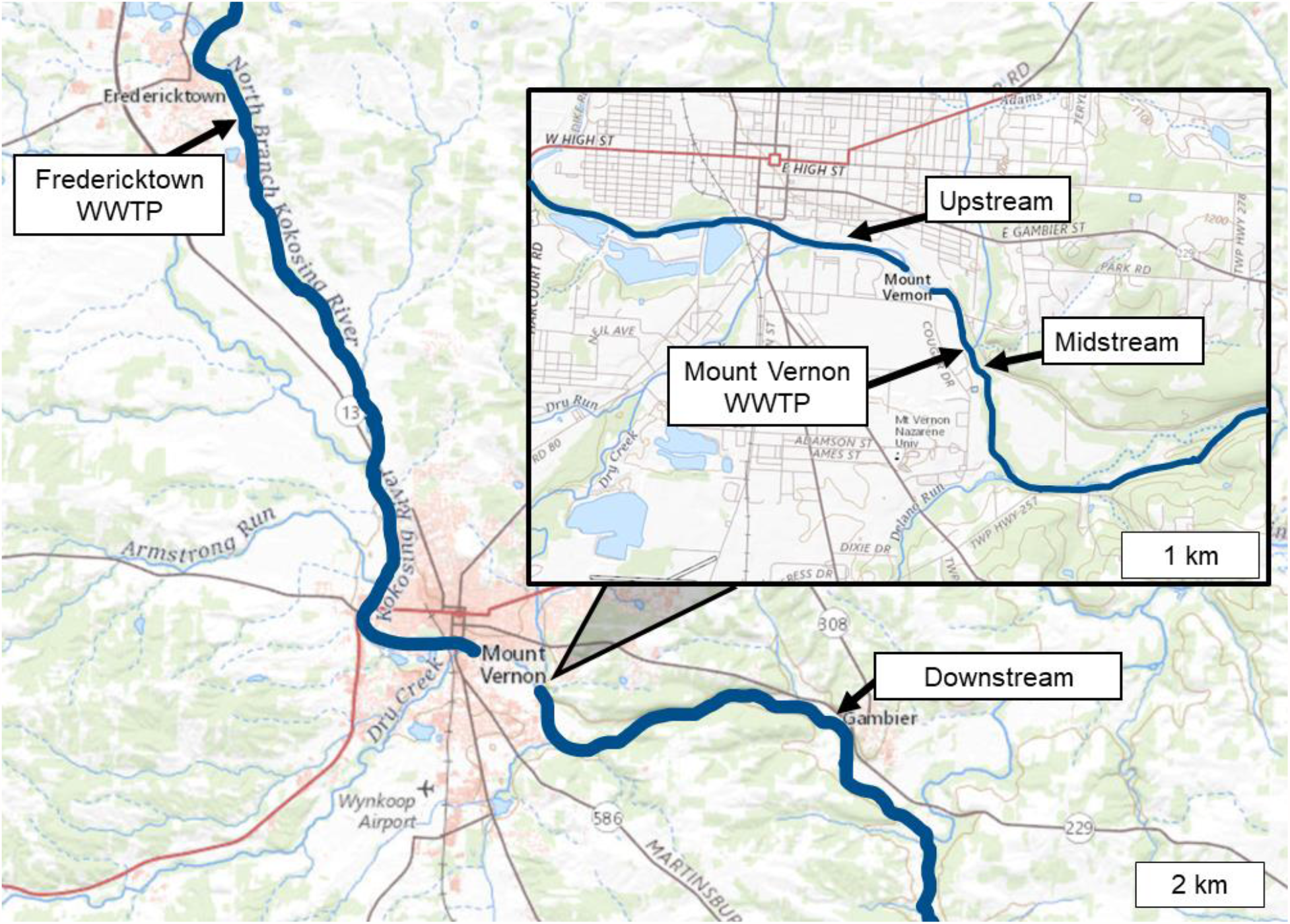
Map of water sampling sites on the Kokosing River. The Upstream site is 1 km upstream of the Mount Vernon City Wastewater Treatment Plant (WWTP). Midstream site is located 9 m downstream of the WWTP. Downstream site is 6 km downstream of the WWTP. Map was generated using the National Wild and Scenic Rivers System (2021).

On each sampling date, the three sites were sampled within a 2-h period. At each site, 400 ml water was collected from the river using a dipper and sealed in sterile WhilrPak® Bags. Within 24 h of sample collection, three 100-ml samples were vacuum-filtered through a sterile 0.22-micron filter, 45 mm in diameter. Filter paper was folded using sterile forceps and deposited in centrifuge tubes which were then frozen at −80°C to preserve microbial DNA. Water pH, conductivity, temperature, and dissolved oxygen (DO) were measured in the field using a Hannah pH/conductivity combination meter and a YSI Pro20 DO meter (Yellow Springs Instruments). Nutrient concentrations were analyzed using collected water samples within 24 h using a portable a Hach® DR900 Multiparameter Portable Colorimeter, including nitrate (NO_3_^−^--N), ammonia (NH_3_-N) and phosphate (PO_4_^−^—P; **Table S1, Supporting Information**).

### DNA isolation and sequencing

Metagenomic DNA was isolated using a ZymoBIOMICS DNA Miniprep Kit. For control samples, 2 μg of the ZymoBIOMICS Microbial Community Standard was processed under the same conditions. This community standard contains defined proportions of ten microbes (5 Gram positive, 3 Gram negative, 2 fungal).

Each filter was cut into small pieces and transferred to a ZR BashingBead Lysis Tube. 650 µl ZymoBIOMICS Lysis Solution was added, and all tubes were processed on a Vortex Genie 2 for 40 min. The remainder of the preparation was performed according to the manufacturer’s protocol. Shotgun sequencing of DNA was performed by Admera Health (www.admerahealth.com). Libraries for sequencing were prepared using Illumina’s Nextera XT DNA Library Preparation Kit, following manufacturer’s instructions. Final libraries were then pooled and sequenced on Illumina HiSeq X sequencer for 150-bp read length in paired-end mode, with an output of 40 million reads per sample.

### ARG marker analysis

Sequence reads were analyzed for ARG marker hits using ShortBRED, a computational pipeline from Huttenhower Biobakery (38). ShortBRED-Identify was used to create a database of short marker peptides specific to ARG protein families compiled from the Comprehensive Antibiotic Desistance database (CARD) (39). From the ARG families, short consensus peptides were identified based on regions of amino-acid sequence identity. To maintain high specificity, the set of peptides was then filtered against the Univeral Protein Database UNIREF90 (https://www.uniprot.org/uniref/) (data accessed October 23, 2019). This database was used to eliminate markers that match sequences outside a specific ARG. One additional marker, ARO_3002930 (vanRO, *Rhodococcus_hoagii*) was removed from the marker set because it lacked specificity. The final list of markers used for our study (ShortBRED-2019) is presented in **Supplemental Table S2**.

The ShortBRED-2019 marker list was used to screen metagenomic reads from each of the three river sites, from six sampling dates (**Supplemental Table S3**). Total read counts per sample were determined using Trimmomatic (40) (**Supplemental Table S4**).

### Taxa profiles

The microbial taxa were profiled using the Huttenhower lab pipeline MetaPhlAn2 (Metagenomic Phylogenetic Analysis) MetaPhlAn2 (41, 42). MetaPhlAn2 assigns metagenomic reads to taxa using a set of clade-specific marker genes identified from approximately 17,000 microbial reference genomes. Taxa were grouped at the levels of phylum, class, order, family, and genus (**Supplemental Table S5**). For control, MetaPhlAn2 was also used to predict the taxa of ZymoBIOMICS Microbial Community Standards that had been prepared concurrently with our experimental samples. For all preparation sets, MetaPhlAn2 consistently predicted the genera of the eight bacterial components and one fungal component of the standard (**Table S7**).

### Data Analysis

To generate an ARG heatmap from the ShortBRED data, we employed R Studio ® 1.3.073. For One-Way ANOVA analysis, we used JMP ® 14.2.0. One-Way ANOVA was used to analyze the significance of ARG and taxa variances among the sites. ARG hits and metadata were correlated by the Spearman rank correlation using R (**Supplemental Table S6**).

### Data submission

For all DNA sequences, FASTQ files were submitted to NCBI, SRA accession number PRJNA706754.

## RESULTS

### ARGs are elevated downstream of the WWTP effluent

We sought to determine how the ARG distribution of the Kokosing River microbiomes was affected by the effluent from the Mount Vernon WWTP. Microbial samples were obtained from three sites on the Kokosing River, designated Upstream (1.0 km upstream of the WWTP), Midstream (120 m below the effluent pipe) and Downstream (6.4 km downstream of the effluent pipe) (**Fig. 1**). From all sites the metagenomic DNA sequences were analyzed for ARG prevalence using the ShortBRED pipeline (38) applied to the CARD database (39). For each marker, the numbers of read hits were summed across all samples and dates, and the markers were ranked according to total hits (**Supplemental Table S3**). Results for the top 60 scoring markers are presented as a heat map (**Fig. 2**).

**Figure 2.**
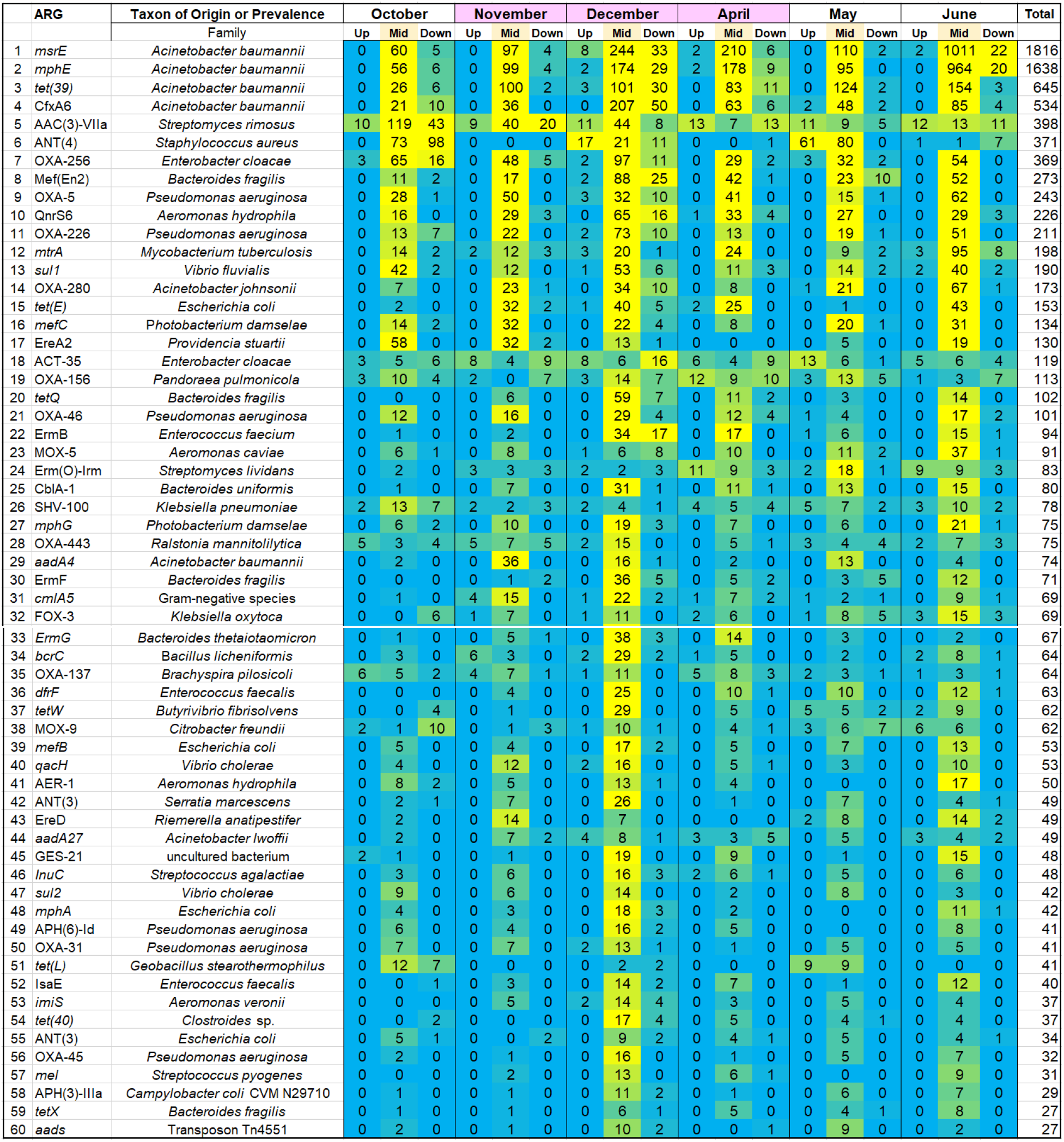
Heatmap of relative abundance of top 60 ARG marker hits. Read hit numbers are ranked in descending order by total hits across samples. Yellow represents highest abundance, cyan represents lowest abundance.

Most of the top-scoring ARGs were elevated in the Midstream samples, compared to samples from either Upstream or Downstream. The elevated ARGs include resistance determinants from several organisms that are of clinical concern. Most striking, six of the abundant ARGs are associated with the ESKAPE pathogen *Acinetobacter baumanii* and related strains: *msrE, mphE, tet(39)*, CfxA6, *oxa280*, and *aadA4* (26–28). The three top-ranked ARGs (*msrE, mphE, tet(39)*) are found together on *A. baumanii* plasmid pS30-1 (34). Overall, the four top *A. baumanii* ARGs account for 37% of the total ARG hits found.

We tested whether the Downstream samples show evidence of carryover from the Midstream site. First, the numbers of ARG hit reads were re-sorted by Midstream site ARG totals (**Table 1**). For each of the top-ranked Midstream ARGs, we present the difference in ARG hits between Upstream and Downstream. The top 27 most abundant Midstream ARGs, including those associated with *Acinetobacter*, all show higher numbers at the Downstream site compared to the Upstream location. In addition, we ranked Midstream ARGs separately for each of the six individual sampling dates and tested the top 20 ARGs for evidence of persistence downstream, using the Wilcoxon signed rank test (**Figure S2)**. Four of the six dates showed significant increase of Midstream ARGs at the Downstream site compared to the Upstream site (P < 0.0083, with Bonferroni correction).

**Table 1.** Top-ranked Midstream ARG hits: Difference in hit numbers between Downstream and Upstream Sample Sites□.

| Mid Rank | ARG Family | Host Organism for ARG (example) | Reads: Down - Up |
| --- | --- | --- | --- |
| 1 | *msrE* | *Acinetobacter baumannii* | 60 |
| 2 | *mphE* | *Acinetobacter baumannii* | 64 |
| 3 | *tet(39)* | *Acinetobacter baumannii* | 51 |
| 4 | CfxA6 | *Acinetobacter baumannii* | 70 |
| 5 | OXA-256 | *Enterobacter cloacae* | 28 |
| 6 | Mef(En2) | *Bacteroides fragilis* | 36 |
| 7 | AAC(3)-VIIa | *Streptomyces rimosus* | 34 |
| 8 | OXA-5 | *Pseudomonas aeruginosa* | 9 |
| 9 | QnrS6 | *Aeromonas hydrophila* | 25 |
| 10 | OXA-226 | *Pseudomonas aeruginosa* | 16 |
| 11 | ANT(4_-)Ib | *Staphylococcus aureus* | 38 |
| 12 | *mtrA* | *Mycobacterium tuberculosis* | 8 |
| 13 | *sul1* | *Vibrio fluvialis* | 12 |
| 14 | OXA-280 | *Acinetobacter johnsonii* | 11 |
| 15 | *tetE* | *Escherichia coli* | 4 |
| 16 | EreA2 | *Providencia stuartii* | 3 |
| 17 | *mefC* | *Photobacterium damsela* | 7 |
| 18 | *tetQ* | *Bacteroides fragilis* | 9 |
| 19 | OXA-46 | *Pseudomonas aeruginosa* | 9 |
| 20 | MOX-5 | *Aeromonas caviae* | 11 |
| 21 | CblA-1 | *Bacteroides uniformis* | 2 |
| 22 | ErmB | *Enterococcus faecium* | 17 |
| 23 | (*aadA4*) | *Acinetobacter baumannii* | 1 |
| 24 | *mphG* | *Photobacterium damsela* | 6 |
| 25 | ErmG | *Bacteroides thetaiotaomicron* | 4 |
| 26 | *dfrF* | *Enterococcus faecalis* | 2 |
| 27 | ErmF | *Bacteroides fragilis* | 14 |

\*ARGs shown represent the top 30 most abundant ARGs from Midstream sites. Values in right- hand column indicate the difference between total Downstream reads and total Upstream reads that match the marker shown.

The overall percentage of reads that matched ARG markers ranged from 0.0015-0.0052% for Midstream samples, and from 0.0002-0.0010% for Upstream and Downstream samples. These numbers indicate roughly 5-fold elevation of ARG hits in the Midstream, compared to the other two sites. We considered the possible effect of sample size, that is, whether the ARG hit numbers reflect the number of reads in our samples (**Supplemental Table S4**). The read counts from individual samples deviated less than 20% from the mean. There was no significant difference in read numbers amongst the three collection sites Upstream, Midstream, and Downstream. Thus, the elevated number of ARGs near the effluent pipe was independent of the number of sequenced reads per sample.

### Taxa profiles associated with WWTP effluent

We investigated whether the elevation of ARGs by the WWTP was associated with specific microbial taxa. The taxa structure of our river metagenomes was determined using the pipeline MetaPhlAn2 (41). The distribution of major bacterial phyla and classes in our samples is shown in **Fig. 3A**, with p-values for Wilcoxon rank sum test (**Fig. 4A**). Reads annotated to the genus *Acinetobacter* showed a striking prevalence in the Midstream, accounting for as high as 30% of predicted organisms (June sample, Midstream); and in some months elevated levels persisted downriver (December and April). By comparison, through ShortBRED, ARGs associated with *A. baumanii* ARGs accounted for 37% of the total ARG hits. This result is striking, since the ShortBRED and MetaPhlAn2 pipelines use very different marker sets (ARGs versus core genome components). Thus the two pipelines offer orthogonal evidence consistent with a high level of multidrug-resistant *A. baumanni* associated with the WWTP plant effluent.

**Figure 3.**
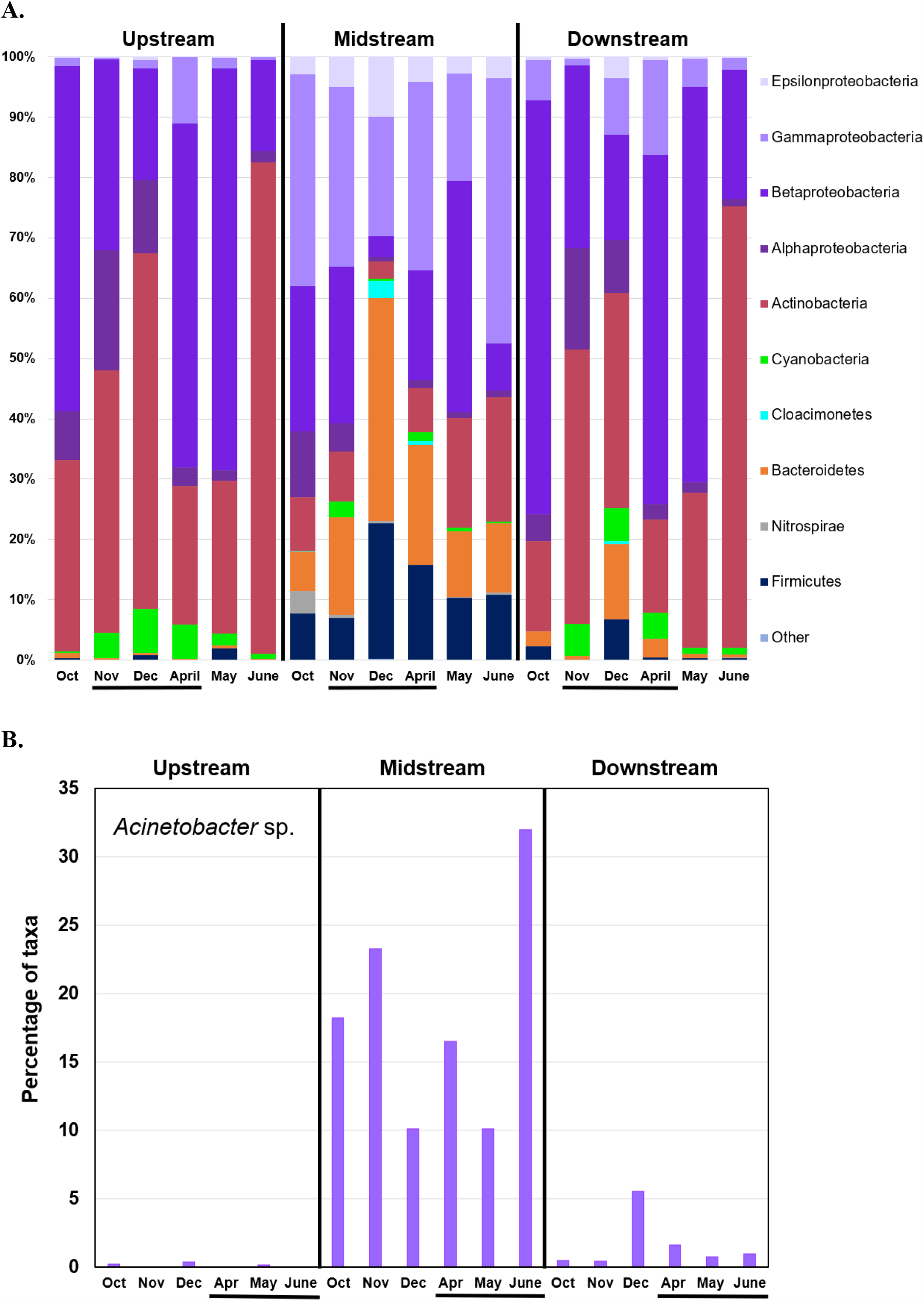
ARGs and bacterial taxa distribution across sampling sites and dates. **A.** Percentages of phyla and of proteobacterial classes predicted by MetaPhlAn2. Taxa with prevalence too small to be quantified were grouped as “Other.” Samples are sorted by site, then by sampling date. Horizontal black bars indicate dates when effluent was unchlorinated. **B.** Percentage of *Acinetobacter* species predicted by MetaPhlAn2.

**Figure 4.**
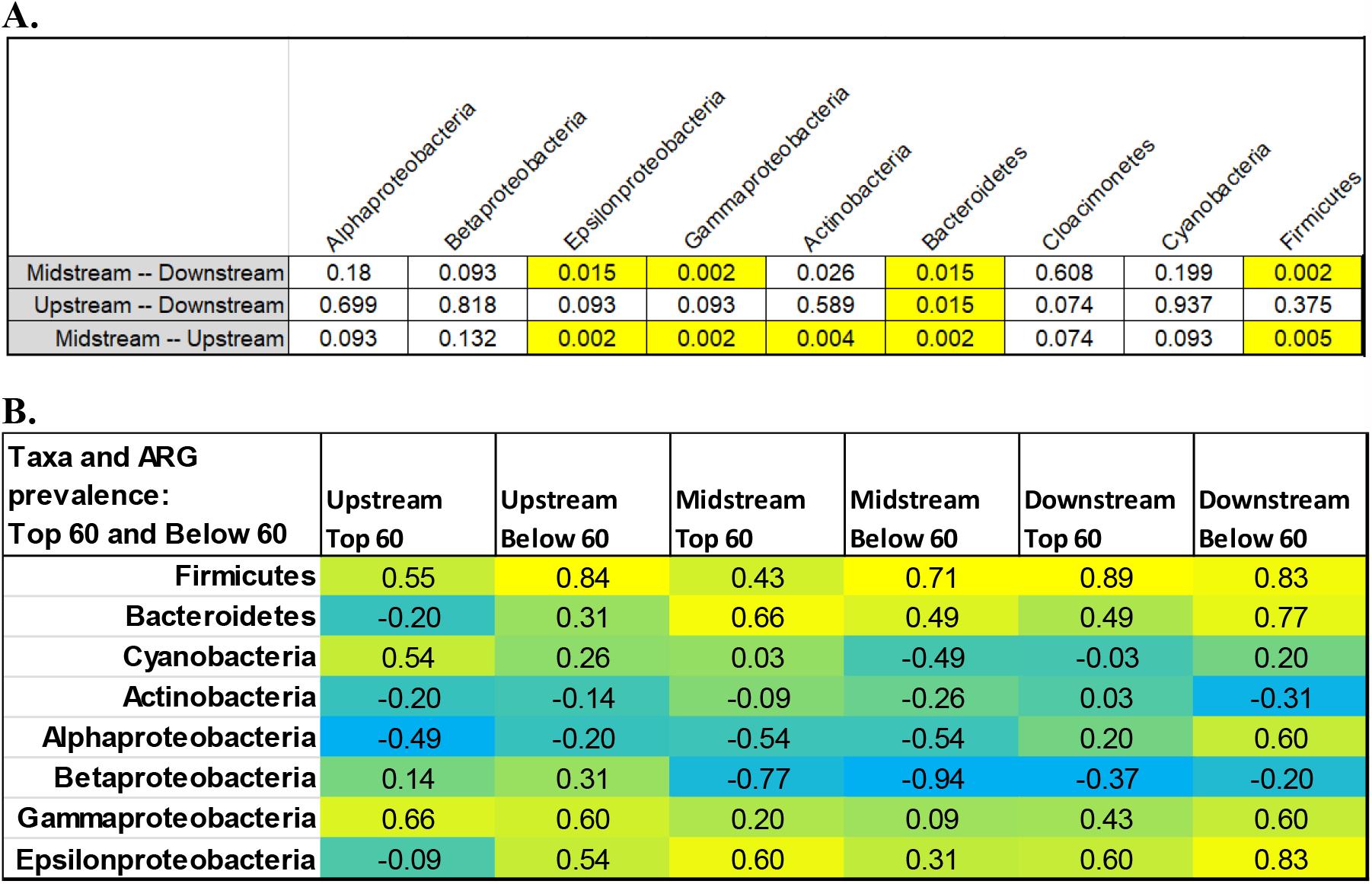
Taxa correlations across river sites. **A**. Wilcoxon rank sum test p-values for comparison of taxa percentages at different sites, grouped across all dates. p-values ≤ 0.02 indicate significant differences (highlighted). **B**. Spearman rank correlations between bacterial taxa and ARG abundance.

The Midstream site showed significantly higher proportions of several major taxa than those Upstream (**Fig. 3A**). The taxa with greater abundance include Bacteroidetes (p=0.002), Epsilonproteobacteria (p=0.002), Gammaproteobacteria (p=0.002) and Firmicutes (p=0.005). Downstream taxa appeared largely similar to those Upstream, with the exception of elevated abundance of Bacteroidetes (p=0.015). These four taxa are consistent with a human fecal source, during the period of effluent chlorination as well as during absence of chlorination. The Upstream and Downstream sites showed higher proportions of Actinobacteria relative to the Midstream. Alphaproteobacteria and Betaproteobacteria showed high prevalence across all three sites. High levels of Actinobacteria and Betaproteobacteria are consistent with metagenomic studies of freshwater oligotrophic lakes and rivers (43).

We considered whether the Midstream elevated ARGs might be associated with bacterial clades that were enriched in Midstream samples. A Spearman rank correlation was performed comparing ARG hits with the major taxa identified (**Fig. 4C**). ARGS were categorized as “Top 60” and “below 60” based on overall rank prevalence (**Fig. 2** and **Supplemental Table S3**). The “Top 60” were those ARG classes showing relative elevation at the Midstream site near the WWTP effluent, whereas ARGs “Below 60” (ranked below the top 60 ARGs) more likely represent autochthonous genes commonly found in a relatively undisturbed river ecosystem. The number of ARG hits at Midstream and Downstream showed a positive correlation with Firmicutes and Epsilonproteobacteria, taxa that might be expected to arise from the WWTP effluent. Negative correlations were seen between ARGs and Betaproteobacteria, which are most likely native to the river.

If the source of “top 60” ARGs is the WWTP, are they carried by the genomes of effluent bacteria, or do they enter the river in the form of environmental DNA? The answer is unclear from our data. However, the occurrence of effluent chlorination (during the months of November, December and April) shows no significant effect on the Midstream taxa profiles (**Fig. 3A**). If live bacteria are responsible for ARGs elevation, significant numbers must be surviving chlorination.

### Nitrate, phosphate and ammonia levels show no correlation with elevated ARGs

The Mount Vernon WWTP effluent commonly includes total suspended solids 1-37 mg/L, phosphorus 2.6-4.1 mg/L, nitrate plus nitrate 5.86-28.9 mg/L, ammonia 0.107-5.77 mg/L (summer), 0.31-10.5 mg/L (winter) (14). Consistent with the above data, our Midstream water samples showed elevated levels of nitrate, phosphate and ammonia relative to the Upstream and Downstream Sites (**Fig. 5** and **Supplemental Table S1**). We therefore looked for possible correlations between water chemistry and ARG prevalence. Spearman rank correlations were performed for ARG levels and various chemical and physical factors (**Supplemental Table S5**). Correlations were run separately for the sums of Top 60 ARG hits and for the sums of Below 60 ARG hits. We hypothesized that the top 60 ARGs are dominated by the WWTP effluent and would therefore show stronger correlations with the Midstream chemistry.

**Figure 5.**
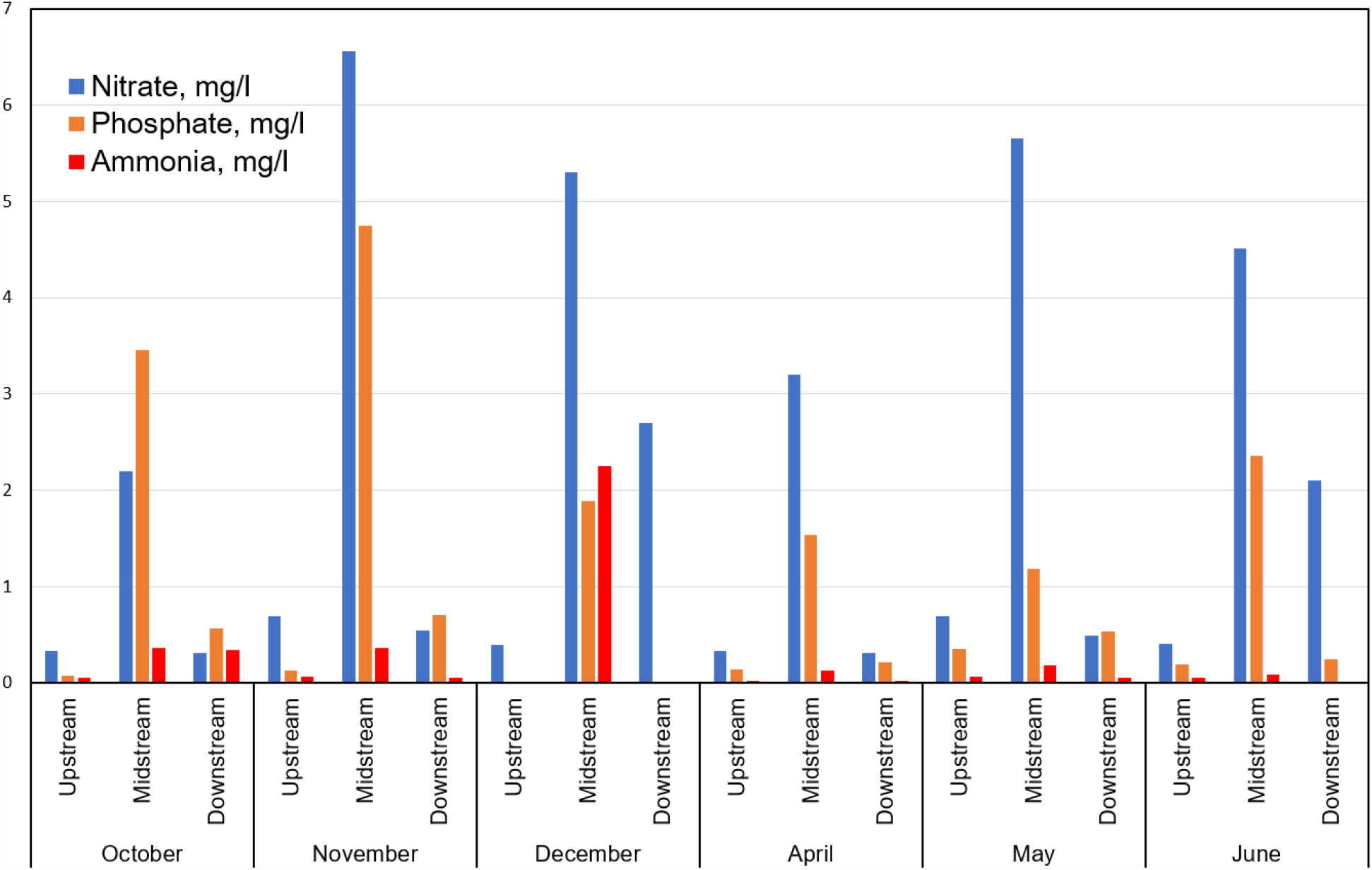
Nitrate, phosphate and ammonia concentrations across sites and dates. Concentrations of nitrate, phosphorus and ammonia were measured at each river site for each month. Full metadata are presented in **Supplemental Table S1**; and metadata correlations with ARG abundance in Supplemental **Table S6**.

In fact, the nitrate, phosphate and ammonia levels showed no consistent correlations with ARGs, either Top 60 or Below 60. This finding suggests that, despite the higher concentration of these nutrients near the effluent pipe, the elevated levels of nitrogen and phosphorus are not correlated with the increased level of ARGs.

The plant effluent typically has a dissolved oxygen content (DO) of 5.3-10.2 mg/L (14). In the Kokosing river samples, we observed DO values ranging from 8.22-12.60 mg/L (Table S1). There was no significant correlation between river DO values and ARG prevalence (Spearman rank correlations, Table S6).

Electrical conductivity (EC) was measured in the Kokosing samples, which has been shown to be an indirect indicator for dissolved organic carbon (DOC) (44–46). Previous studies find connections between DOC and ARG abundance (47, 48). Over the courrse of our study, EC values ranged from 500-890 µS/cm (Table S1) but no significant correlation was found with site location or season, nor with ARG levels (Table S6).

### ARG numbers increased with pH and temperature

The strongest strongest correlations we saw between ARGs and water chemistry were for pH and temperature (**Supplemental Table S5**). The range of pH values observed was pH 7.19-8.55 (**Supplemental Table S1**). At Midstream and Downstream sites, pH showed positive correlations with ARG hits, particularly the Below 60 ARGs. These results suggest the possibility that low pH might select against ARGs that commonly occur in metagenomes of the undisturbed river. In laboratory evolution experiments on *Escherichia coli*, low pH and membrane-permeant aromatic acids select for loss of ARGs and ARG regulators (49, 50).

Temperature showed a strongly negative correlation with ARG levels, particularly those Below 60. This finding suggests the possibility of high-temperature selection against ARGs commonly found in the river community.

## DISCUSSION

Past studies have investigated ARGs in urban waterways, but there has been relatively little research on the occurrence of ARGs in rural watersheds characterized by low human population density and agricultural land use. In addition, few studies have focused on rivers that are considered to be of exceptional quality, such as the Kokosing River investigated here. Forty-seven miles of the river are designated “scenic” by the state, and the river attracts members of the public for fishing, birding and canoeing. Nonetheless portions of the river are impacted by livestock and agriculture, as well as pollution from a residential lakeside development (14). In 2007, portions of the watershed were reported to be impacted by gravel mining, erosion, and conversion to row crops.

Despite the overall high water quality of this river system, and the relatively small contributions of WWTP effluent to stream discharge, we found substantially higher ARG abundance downriver of a WWTP compared to the more agricultural portions of the watershed that lie upstream. we found substantially higher ARG abundance downriver of a WWTP compared to the more agricultural portions of the watershed that lie upstream. The WWTP influx inputs a few percent of the total river flow rate (Supplementary Table S1). Thus, a relatively small city WWTP (catchment population 17,000) may foster the spread of ARGs in a river that is in excellent ecological condition, as has been shown for anthropogenic contaminants in large, urban centers (see for example (51)).

The footprint of the WWTP effluent release was evident across our data, including shifts in ARG prevalence (**Fig. 2, Table 1**), microbial community taxa distribution (**Fig. 3**), and chemical indicators (**Fig. 5**). The top three ARGs for ShortBRED markers ranked in our metagenomes are known to occur together on *A. baumanii* plasmid pS30-1 (34). In addition, the MetaPhlAn2 taxonomic pipeline, with completely different markers, found high prevalence of *A. baumanii* near the WWTP effluent (**Fig. 3B**). It is possible that the multidrug-resistant *A. baumanii* actually comes from the WWTP. Wastewater treatment is known to increase the prevalence of multidrug resistance in *A. baumanii* from influent to the final effluent (12).

An effect of the wastewater effluent could be to increase community exposure to drug-resistant strains of this ESKAPE pathogen. It is also possible that the ARGs associated with *A. baumanii* have been acquired by other members of the native river microbial community. Nevertheless, the possibility of *A. baumanii* contamination should be followed up by further studies. *Acinetobacter* species of concern are emerging worldwide, especially in warmer climates; and their prevalence likely will increase with climate change (52–54). River levels of *Acinetobacter* species can be examined by targeted metagenomic analysis (55), amplicon assessment (56) and culture-based methods (57).

Rural rivers have substantial economic and cultural significance for local human communities. Nevertheless, the public is rarely aware of the potential impact of WWTP ARG exposure, with the common presence of WWTP plants along rural rivers. For example, 25 km upstream of the Mount Vernon plant is the Fredericktown WWTP; and just downstream of our sampled sites in Gambier, another small WWTP releases effluent to the Kokosing. Further downstream from Gambier (20 km) lies the Danville WWTP (design flow 0.20 MGD).

We found evidence that detectable levels of ARGs persist in the river microbial community at least several kilometers past the effluent pipe. The Downstream site exhibited higher ARG counts than Upstream for the top 27 ARGs elevated at Midstream (**Table 1**). Thus, WWTP-associated ARGs persist and are transported in the environment at least 6.4 km downstream. Most of these ARGs are found in multiple species and may be transmitted by mobile elements (39). These ARGs might become established in the river microbial resistome and could propagate to pathogenic bacteria in the future, posing a risk to human health.

The WWTP-proximal site also showed substantial alteration of overall taxa distributions, such as increased prevalence of Bacteroidetes and Firmicutes (Fig. 4), findings that are consistent with previous study (35). The increase in Bacteroidetes persisted 6.4 km downstream. In addition, the WWTP-proximal site showed depletion of Actinobacteria, although the levels of this river group recovered downstream.

There is need for future investigation regarding efficient methods of ARG control from WWTP in freshwater systems (59). In addition, the public should be more aware of the entry of wastewater into recreational waterways. Better awareness of the consequences of WWTP-effluent release into rivers will improve our ability to sustain healthy microbial communities in our freshwater systems.

## Acknowledgements

This work was supported by the National Science Foundation award MCB-1923077. We acknowledge contributions of students in the Kenyon College course BIOL 239 Experimental Microbiology.

## Notes

### Competing Interest Statement

The authors have declared no competing interest.

### Summary of Updates

Added data on WWTP and Kokosing watershed. Corrected sampling site locations.

